# Non-contact optical imaging of tissue deformation enables *in vivo* cardiovascular monitoring

**DOI:** 10.64898/2026.09.04.749131

**Authors:** Austen T. Lefebvre, Nicole E. Steiner, Siyu Wang, Carissa L. Rodriguez, Eyal Bar-Kochba, Yanrong Shi, Sung-Min Cho, David W. Blodgett, Marek Mirski

**Author notes:** Address all correspondence to Austen Lefebvre. These authors contributed equally to this work (Austen T. Lefebvre, Nicole E. Steiner).

## Abstract

**Significance:** Continuous, non-invasive monitoring of cardiovascular physiology provides critical information for clinical decision-making and patient care. Emerging optical sensors enable non-contact measurement of physiological parameters such as heart rate and respiratory rate, but current methods are limited in their ability to capture spatially resolved physiological waveforms.

**Aim:** We aimed to extend the capabilities of non-contact cardiovascular monitoring by using an imaging-based approach from which spatially resolved physiological waveforms can be extracted and cardiovascular biomarkers can be derived *in vivo*.

**Approach:** We implemented a digital holographic imaging sensor to continuously measure calibrated tissue motion for *in vivo* assessment of cardiovascular biomarkers in six adult male Sprague-Dawley rats, with validation against electrocardiographic and invasive arterial blood pressure measurements.

**Results:** Heart rate and pulse arrival time-derived pulse wave velocity calculated from digital holographic imaging demonstrate strong agreement with reference-derived metrics (concordance correlation coefficient ≥ 0.98). Heart rate variability shows moderate agreement with reference-derived metrics (concordance correlation coefficient ≥ 0.59).

**Conclusions:** Digital holographic imaging is a promising modality for non-contact cardiovascular monitoring, allowing for continuous measurement of tissue motion that enables assessment of key cardiovascular biomarkers.

## 1. Introduction

Continuous physiologic monitoring is an essential component of clinical care, providing real-time insight for diagnoses and treatment of patients. Non-invasive and non-contact sensors are capable of recording a wide range of clinically relevant, continuous physiologic signals that help guide clinical decision making and management of patients [1]. Of particular interest is tracking cardiovascular function, where real-time monitoring helps guide management in diseases such as sepsis and cardiogenic shock [2].

Non-invasive measures of tissue motion dynamics can provide clinically relevant insight into cardiac physiology by quantifying changes in the macroballistic motion of tissue driven by the cardiac cycle [3–5]. Ballistocardiography (BCG) exploits this signature for monitoring cardiovascular physiology by recording the whole body motion that occurs due to cardiac ejection of blood and vascular flow [5]. Recordings of these mechanically-derived waveforms have the potential to non-invasively inform cardiac output [6] and blood pressure [4]. BCG has also shown promise in the assessment and management of patients with heart failure [7]. Seismocardiography (SCG) is a similar approach that records chest wall vibrations induced by the motion of the heart to inform cardiovascular health [8]. SCG has shown promise in measuring cardiovascular metrics such as heart rate variability via wearable sensors on the thorax [8–10].

BCG and SCG have demonstrated that non-invasive measurements of the motion of tissue can be used to derive clinically relevant cardiac information across a range of approaches and sensors, but they largely require contact with the tissue [6–11]. Laser Doppler vibrometry (LDV) has emerged as a promising non-contact approach to measuring tissue motion. LDV is an optical technique that measures the Doppler phase and frequency shift imparted on a laser beam by vibrating objects [12]. Similar to BCG and SCG recordings, LDV has been shown to provide insight into cardiovascular physiology by measuring displacements from arterial pulsations for monitoring heart rate as well as vascular health [13,14]. Two challenges with using LDV include its sensitivity to signal drop-out due to random changes in speckle [12] and the need for precision pointing of the instrument to ensure optimum placement of the recording location.

Building on our previous work demonstrating that a full-field digital holographic imaging (DHI) system can measure nanometer-scale tissue deformation associated with neural activity *in vivo* [15], we hypothesize that this approach can be extended to cardiovascular monitoring. Compared with single-point LDV, this full-field interferometric approach should reduce susceptibility to speckle-related signal dropout and the need for precision pointing while retaining the sensitivity required for accurate non-contact cardiovascular recordings. To that end, we use our DHI system to demonstrate that tissue deformation recorded at high spatiotemporal resolution enables non-contact characterization of cardiovascular physiology.

## 2. Methods

### 2.1 Digital holographic imaging system overview

Optical data were acquired with a DHI system capable of recording nanometer-scale full-field, time-dependent out-of-plane tissue motion. Utilizing an off-axis holographic configuration, light scattered from an object (i.e., tissue) is mixed with a reference beam at the focal plane array (FPA) forming a hologram, which is digitally processed to compute tissue velocity.

The DHI system operates in reflection geometry using a 1320 nm, long coherence length laser (QPhotonics Model #QDFBSS-1320-20) that is split into a reference and sample arm. The sample arm is expanded into a 3-mm diameter collimated beam that illuminates the sample. The sample arm illuminates the tissue at a skin-safe operating power, and the scattered light is collected and relayed through a 1:1 telecentric lens system, forming an image 75 mm in front of a short-wave infrared camera (C-RED 3 Axiom Optics). A conventional off-axis hologram is then formed after mixing the scattered sample light with the reference arm using a beam splitter. For this study, the camera acquired holograms with a FPA size of 64 × 64 pixels at a frame rate of 9,596 Hz and exposure time of 90 µs. The reconstructed image spatial resolution, calculated based on the 15 µm pixel pitch and distance between the FPA and image plane, was 120 μm. The system layout and processing approach are detailed in Figure 1.

**Figure 1.**
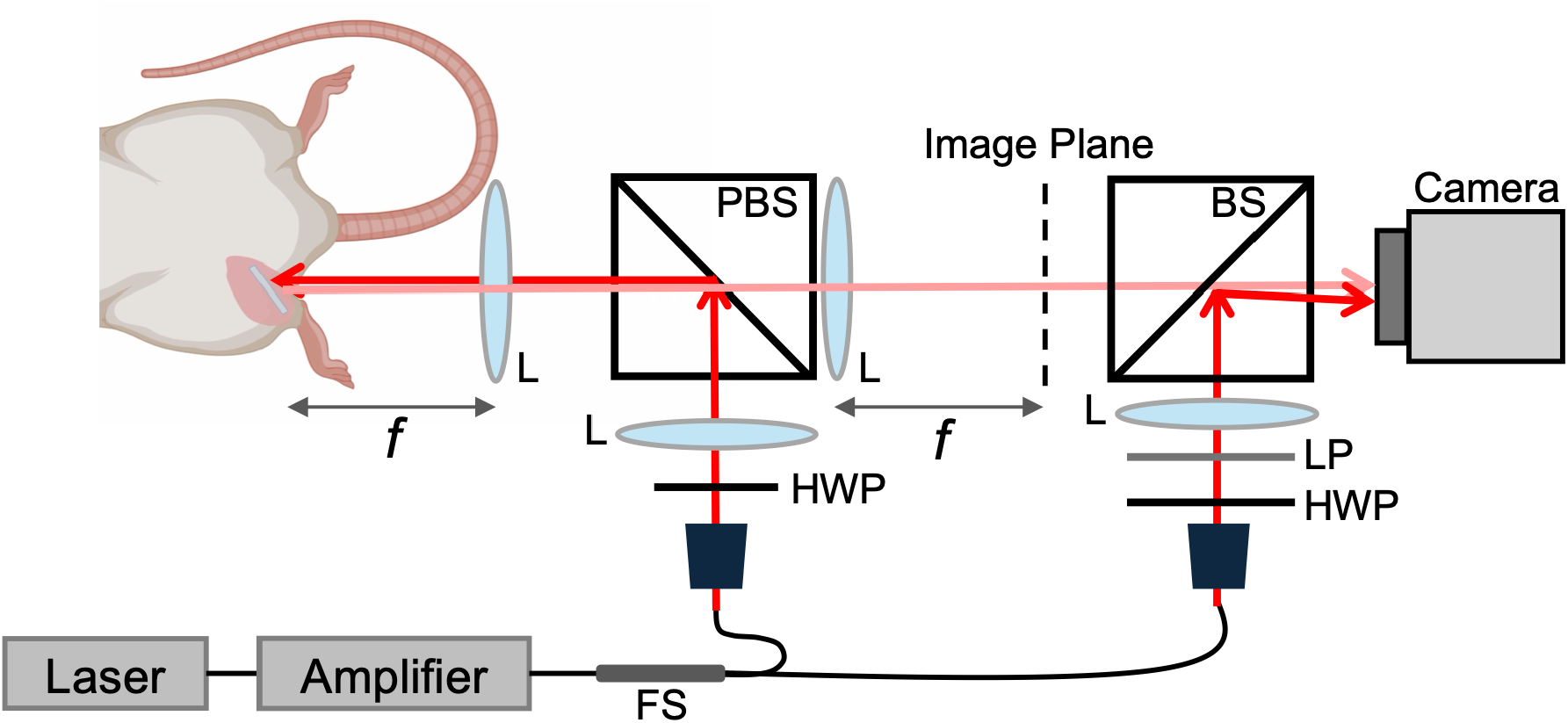
DHI system layout. PBS: polarizing beam splitter, BS: beam splitter, LP: linear polarizer, HWP: half-wave plate, L: lens, FS: fiber splitter, *f*: focal distance. The laser is split between an object and reference arm. Collimated light is directed to the sample via a PBS. Light scattered back from the sample is then collected through the PBS and mixed with the reference arm to form an off-axis hologram at the camera. Part of the schematic created in BioRender. Lefebvre, A. (2025) https://BioRender.com/p6vo82j

### 2.2 Deriving tissue velocity and displacement

Deriving tissue velocity from an acquired stack of holograms involves several steps [15]. First, the hologram *H* is converted to k-space with an FFT, allowing for high-pass spatial filtering of the DC term. Then the Fresnel transform step involves applying an IFFT to go back into hologram space, applying a propagator term to propagate the wavefront to the depth of the target, and then applying an FFT to generate a complex image of the target. The resulting image stack *I* contains magnitude and phase information about the target at each time step. The differential phase for each pixel is then computed by taking the complex cross-correlation between successive frames, using the following equation where x and t denote the spatial and time dimension, respectively, and * is the complex conjugate operation:

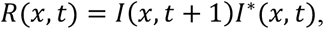

The average differential phase is then spatially averaged over an identified region-of-interest (ROI) as follows:

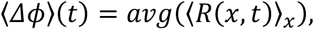

where ⟨.⟩_x_ is the independently averaged real and imaginary components of *R*. Finally, velocity is derived by scaling ⟨Δ*ϕ*⟩ by the wavelength (λ) of the laser and the sampling interval (Δt), as described below. Displacement is derived by simply integrating the velocity signal.

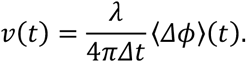

ROIs were manually selected in each data collection to encompass the tissue expected to produce the strongest physiological signal over a 5 × 5-pixel area. Specifically for this study, ROI placement targeted tissue around the femoral artery, and inspection of the resulting waveform was used to ensure adequate signal quality. Figure 2 details the utility of ROI selection and demonstrates the spatial variability of the signal across the field of view.

**Figure 2.**
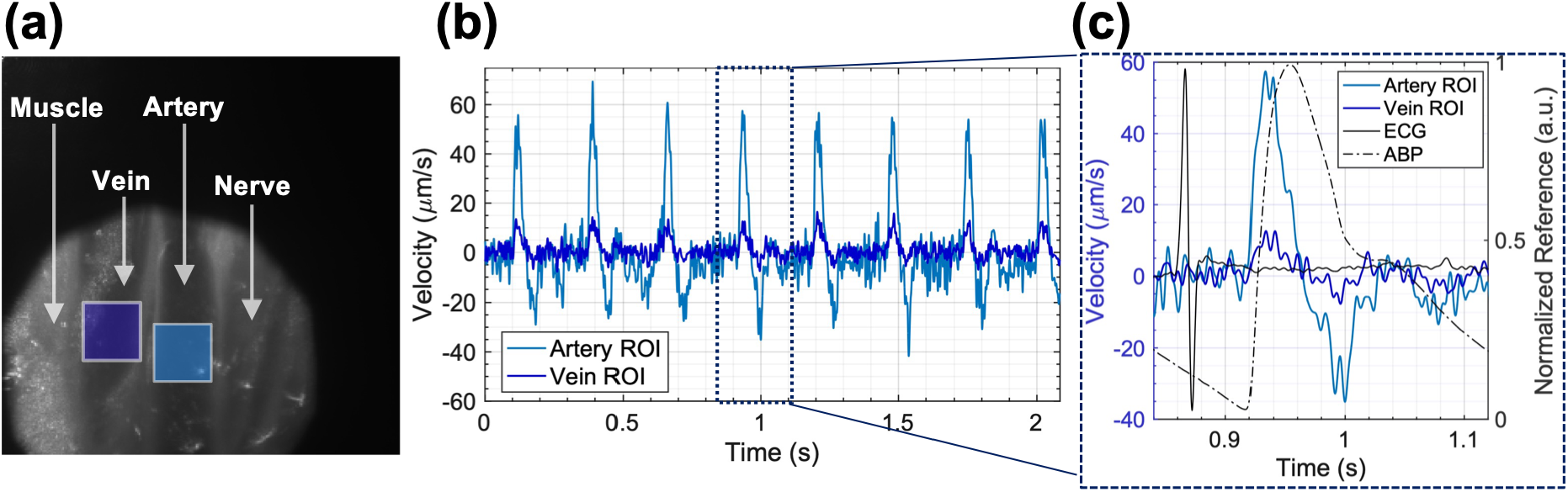
Impact of region-of-interest selection. (a) A high spatial resolution average 1-frame difference image of the DHI sensing region reveals the femoral vein, artery, and nerve of the proximal femoral neurovascular bundle, as well as the adjacent medial thigh musculature. The dark blue and light blue boxes indicate ROIs positioned over the femoral vein and artery, respectively. (b) Velocity waveforms derived from the venous and arterial ROIs identified in panel (a). (c) Expanded view of the single cardiac cycle boxed in panel (b), with overlaid normalized reference electrocardiogram (ECG) and arterial blood pressure (ABP) waveforms. The image in panel (a) was generated from 1,000 consecutive frames (512 × 512 pixels) acquired at 697 Hz; the waveforms in panels (b) and (c) were derived from frames acquired at 64 × 64 pixels and 9,596 Hz using 5 × 5-pixel ROIs.

All data processing and statistical analyses were performed using MATLAB 2024b (MathWorks, Natick, MA, USA).

### 2.3 Animal preparation

All procedures were approved by the Johns Hopkins University Animal Care and Use Committee (ACUC). Male Sprague–Dawley rats (CD® IGS, 350–450 g; Charles River Laboratories, Wilmington, MA) were induced with 3% isoflurane in 100% oxygen delivered by face mask. Anesthesia was subsequently maintained via face mask with 1.0–1.5% isoflurane in an oxygen/air mixture adjusted to 50% oxygen at a total flow rate of 1 L/min, with the isoflurane concentration adjusted according to anesthetic depth. Intraperitoneal (IP) ketamine (25 mg/kg) was administered as supplemental anesthesia, with additional doses administered as needed based on changes in heart rate and blood pressure. Local anesthetic infiltration with 0.2% bupivacaine was administered at surgical sites. A heating pad (RightTemp, Kent Scientific Corporation) was placed beneath the rat to maintain body temperature at 37 °C during the experiment.

The rat was placed supine, and the medial thigh was shaved and disinfected. Subcutaneous tissue was bluntly dissected until the neurovascular bundle (femoral artery, vein, and nerve) was exposed. The left femoral artery and vein were carefully separated from the nerve using fine forceps. The Mikro-Tip catheter transducer (Millar, Inc., SPR-407 2F, Houston, TX) was inserted approximately 20 mm into the lumen of the left femoral artery and connected to a bridge amplifier (ADInstruments Bridge Amp, FE221) for arterial blood pressure recording. A pre-flushed PE-50 catheter containing heparinized saline (100 U/mL) was inserted into the left femoral vein for intravenous drug infusion. The right femoral artery and vein were exposed at a similar anatomical level to the Mikro-Tip catheter transducer on the contralateral side. The DHI measurement sites and Mikro-Tip sensor location were marked to measure the respective distances from the aortic arch.

Additional physiologic measures were continuously recorded, including oxygen saturation (SpO_2_) using a rat paw sensor placed on the right hindpaw (PhysioSuite, Kent Scientific Corporation) and a 3-lead electrocardiogram (ECG) recorded from needle electrodes placed in the left and right forepaws and right hindpaw using a bioamplifier (ADInstruments Quad Bio Amp, FE234). All physiologic signals and camera triggers were acquired with a PowerLab 16/35 data acquisition system (ADInstruments, PL3516) and LabChart 8.1.2. The physiologic signals were recorded at a sampling rate of 4 kHz and camera triggers at 40 kHz. Camera frame timing was aligned with physiologic waveforms via the simultaneously recorded camera trigger channel.

### 2.4 Heart rate and rhythm

Heart rate and rhythm describe the frequency and pattern of cardiac contraction and arise from the intrinsic electrical activity of the heart’s conduction system. This intrinsic activity is continuously modulated by the autonomic nervous system, and heart rate and rhythm reflect the health of the heart as well as autonomic regulation. Therefore, they are essential clinical measures of a patient’s physiologic state, informing the detection and prognosis across a range of pathologic conditions from sepsis and shock to acute coronary syndrome, and serving as established predictors of clinical deterioration and mortality [1].

#### 2.4.1 Data acquisition and processing of arrhythmia

Characterization of tissue velocity during arrhythmias was accomplished with rapid administration of intravenous adenosine, an endogenous purine nucleoside that transiently slows heart rate and atrioventricular (AV) nodal conduction [16,17]. This can result in bradycardia and AV block that may progress to complete AV block. Following collection of baseline data, a rapid intravenous injection of adenosine (2 mg/kg) was administered via a left femoral venous catheter to induce a transient bradyarrhythmia and AV heart block. Optical data were recorded at two locations over the right femoral artery: one over the exposed proximal right femoral artery at approximately the same anatomical level as the Mikro-Tip pressure transducer in the contralateral left femoral artery, and a second, slightly more proximal transcutaneous location where overlying skin and soft tissue remained intact. For rhythm tracking, DHI-derived velocity and ECG signals were low-pass filtered using a sixth-order Butterworth filter with a cutoff frequency of 150 Hz. Around the time of adenosine administration, peaks were manually identified in each signal to track beat-to-beat intervals measured by both modalities.

### 2.5 Heart rate variability and associated metrics

Heart rate variability (HRV) describes variations in the time intervals between consecutive heartbeats. It is indicative of functioning of the autonomic nervous system, which regulates critical physiological processes like blood pressure, digestion, and respiration [18]. HRV has been shown to be related to a multitude of clinically relevant states in humans, including but not limited to: inflammation, infection, drug use, anxiety disorders, and genetic factors [19, 20].

HRV may be computed across different timescales ranging from long (hours) to ultra-short (<5 min) and may be derived using time-domain, frequency-domain, or non-linear calculations [18]. For this study, timescales were adapted to be appropriate for rodent cardiac physiology, and, as such, a shorter timescale (60 s) for deriving time-domain features was utilized. In particular, the commonly used standard deviation of RR intervals (SDRR) and root mean square of successive differences (RMSSD) measures were computed for comparison between ECG-derived R intervals and velocity-derived peak intervals, both referred to as inter-beat intervals. Mean heart rate (HR) is also computed using the average inter-beat interval. Calculations used to derive each of these measures are described in Table 1, where *I* represents a vector of N inter-beat intervals and *I*_*i*_ represents the i^th^ element of the vector.

**Table 1.** HR and HRV metric computations.

| HRV Metric | Equation |
| --- | --- |
| Mean HR | $\frac{60}{\frac{1}{N} \sum_{i=1}^N I_i}$ |
| SDRR | $\sqrt{\frac{1}{N-1} \sum_{i=1}^N (I_i - \bar{I})^2}$ |
| RMSSD | $\sqrt{\frac{1}{N-1} \sum_{i=1}^{N-1} (I_{i+1} - I_i)^2}$ |

#### 2.5.1 Data acquisition and processing of HRV

Each collection was windowed into non-overlapping 60 s windows. In order to compute HRV metrics, R peaks and corresponding peaks were detected in the ECG and velocity signals, respectively, for each window. Mean HR, SDRR, and RMSSD were then derived from the resulting intervals for each modality.

For R peak detection, ECG signals were low-pass filtered using a sixth-order Butterworth filter with a cutoff frequency of 150 Hz. For optical peak detection, velocity signals were bandpass-filtered using a sixth-order Butterworth filter with a low-pass cutoff frequency of 50 Hz to remove high-frequency noise and a high-pass cutoff frequency of 2 Hz to mitigate respiration contributions. Peaks were then detected using MATLAB’s findpeaks function, with the minimum timing distance set at 100 ms and the minimum peak magnitude set to be 3 standard deviations of the respective waveform. Using the detected peak timing, intervals were computed as the time difference between successive peaks. Outlier intervals were identified as any intervals that exceeded the median interval by more than 3 median absolute deviations and removed. Finally, HRV metrics were computed across all remaining intervals for comparison. Representative ECG and velocity waveforms with identified R and corresponding peaks in the velocity waveforms, which will be referred to as PV (peak velocity), alongside a comparison of the derived RR intervals over time and across the entire data collection are shown in Figure 3.

**Figure 3.**
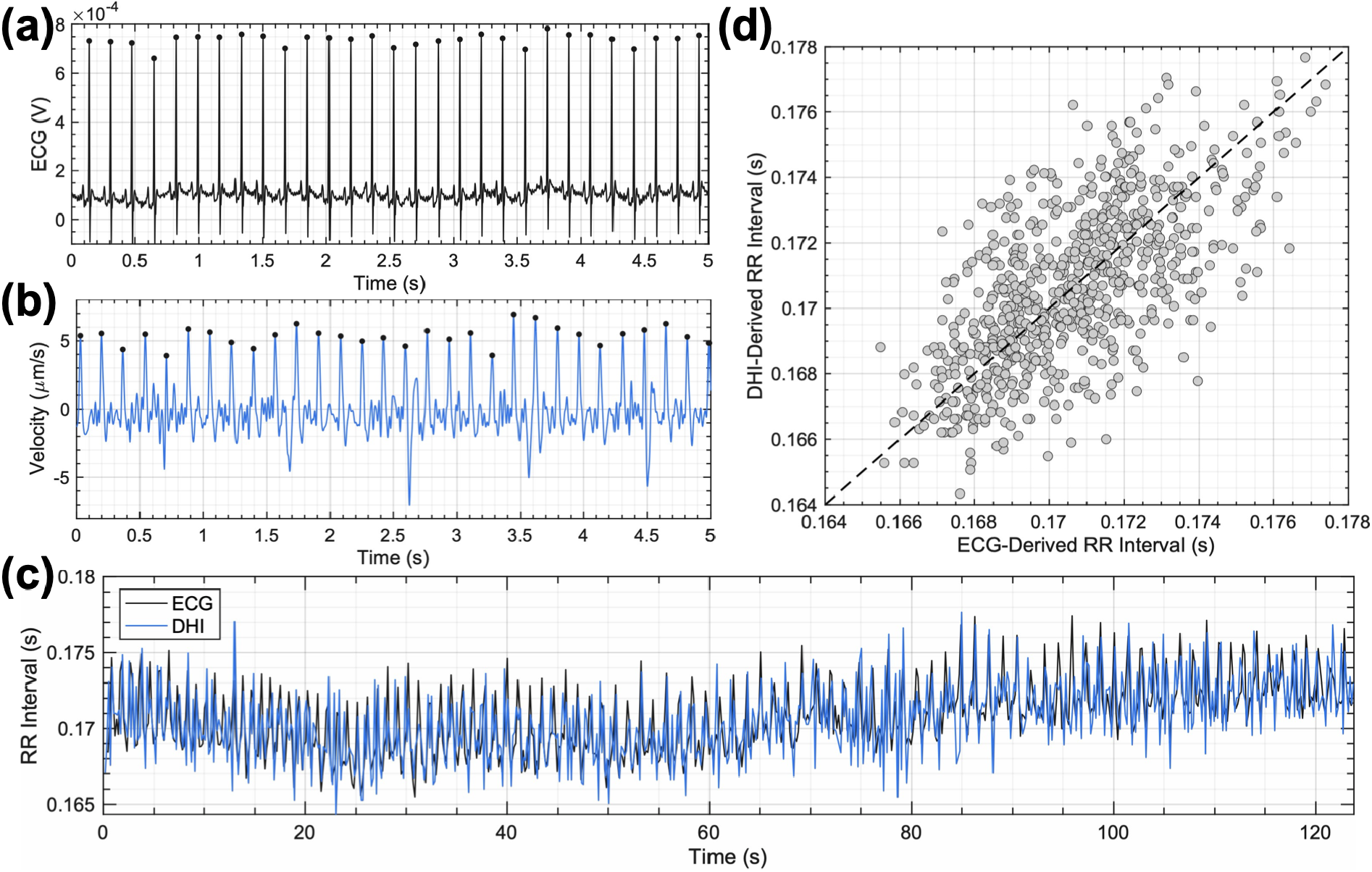
Representative R and PV peak detection and comparison of resulting RR intervals. (a) ECG waveform with overlaid detected R peaks. (b) DHI-derived velocity waveform with overlaid detected PV peaks. (c) RR intervals detected in ECG (gray) and DHI (blue) waveforms over the course of one data collection. (d) Scatter plot comparing RR intervals computed from ECG versus from DHI for the same data collection.

### 2.6 Pulse arrival time and pulse wave velocity

Pulse arrival time (PAT) refers to the time interval between cardiac electrical activation, identified by the ECG R wave, and arrival of the resulting arterial pulse at a peripheral measurement site. PAT includes both the pre-ejection period and the subsequent vascular pulse transit time. Pulse wave velocity (PWV) quantifies the propagation speed of the arterial pulse wave and is conventionally determined from vascular transit time and propagation distance. In this study, PWV was estimated from PAT and the measured arterial path length *[21]*. Because pulse wave propagation is influenced by vascular compliance, arterial stiffness, and blood pressure, both PAT and PWV provide complementary dynamic measures of cardiovascular state [22–25].

#### 2.6.1 Data acquisition and processing of PAT and PWV

DHI-derived tissue displacement signals and invasive arterial blood pressure (ABP) signals were simultaneously recorded from each side of the exposed femoral arteries. For each animal, three anatomical distances were measured: the distance from the aortic arch to the aortic bifurcation (d0), the distance from the aortic bifurcation to the invasive ABP sensor (d1), and the distance from the aortic bifurcation to the DHI measurement site (d2), as shown in Figure 4a. To preserve systolic and diastolic waveform morphology, both ABP and tissue displacement signals were bandpass filtered from 0.5 to 35 Hz using a sixth-order Butterworth filter.

**Figure 4.**
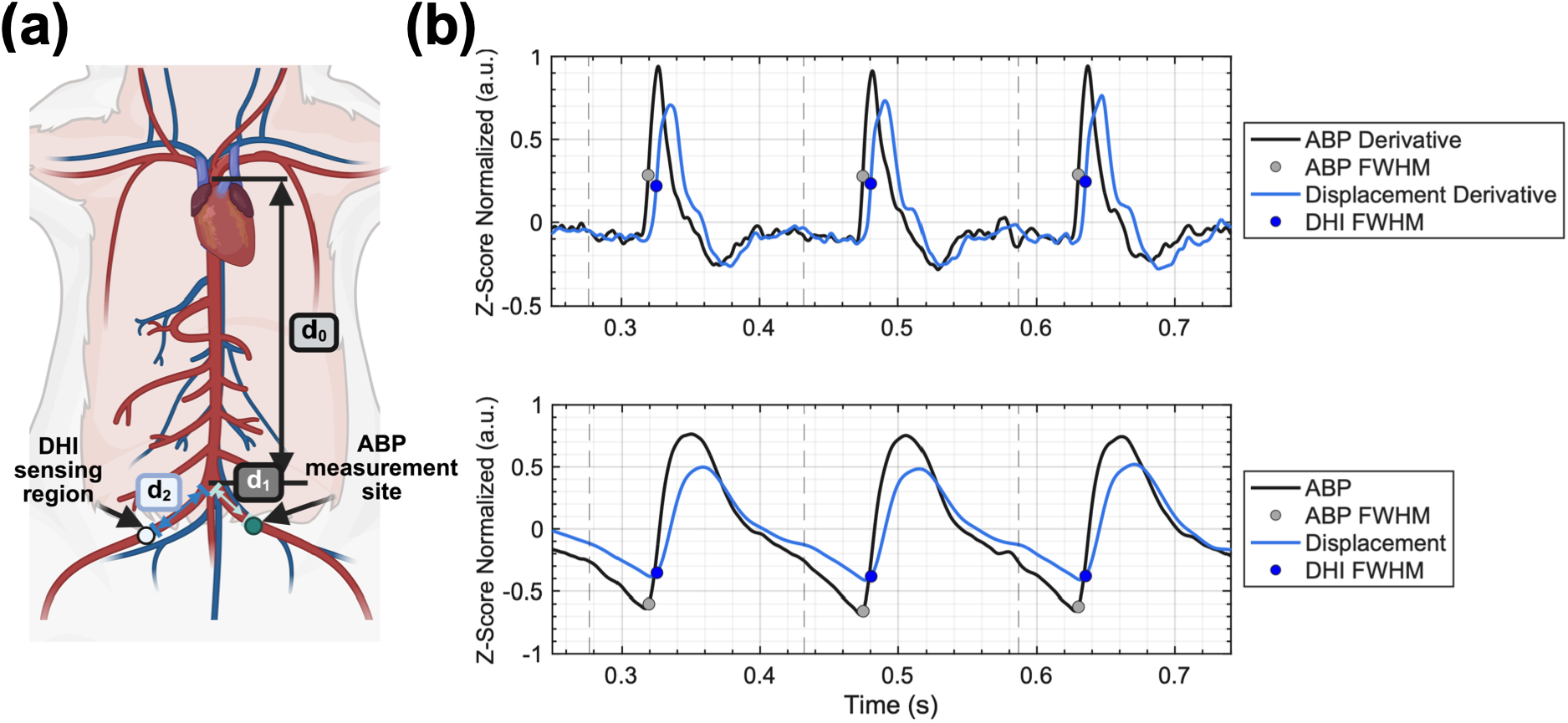
Approach to deriving pulse arrival time (PAT) and PAT-derived pulse wave velocity (PWV). (a) Anatomical distance measurements used to calculate PAT-derived PWV. The propagation distance from the aortic arch to the invasive arterial blood pressure sensor was defined as d0 + d1, and the propagation distance from the aortic arch to the DHI measurement site was defined as d0 + d2. Part of the schematic created in BioRender. Thakor, N. (2026) https://BioRender.com/mibuskz. (b) Representative normalized ABP and DHI-derived waveforms with corresponding first derivatives. The rising-edge timing marker was defined as the time point preceding the derivative peak at which the derivative waveform reached 50% of its maximum value. Dashed vertical lines indicate the R-peak detection times, and PAT was calculated as the time from each R-peak detection to the FWHM-based rising-edge detection time.

The rising edges of the ABP waveform and the tissue displacement waveform were used as pulse arrival markers. A full width at half-maximum (FWHM)-based approach was used to identify the rising edge of each signal [26–29]. Briefly, the first derivative of each waveform was calculated, and the derivative peak was identified using MATLAB’s findpeaks with the minimum timing distance set at 100 ms and the minimum peak magnitude set to be 3 standard deviations of the respective waveform. The rising-edge time point was defined as the time point preceding the derivative peak at which the derivative waveform reached 50% of its corresponding maximum value.

PAT was calculated on a beat-to-beat basis as the temporal difference between the R-peak of the simultaneously recorded ECG and the corresponding rising edge of either the ABP waveform or the tissue displacement waveform. PWV was then estimated by dividing the propagation distance by the corresponding PAT. For invasive ABP sensor measurements, the propagation distance was defined as d0 + d1 (distance from the aortic arch to the arterial pressure catheter), whereas for DHI measurements, it was defined as d0 + d2 (distance from the aortic arch to the center of the DHI measurement site) [26,30]. Differences in PAT between the two modalities were expected because the arterial catheter and DHI measurement sites were located at different distances from the aortic arch. Intervals exceeding the median by more than three MADs were classified as outliers and excluded from subsequent performance metric analyses [31].

### 2.7 HRV and PWV agreement analysis and performance

Agreement of HRV and PWV between DHI and corresponding ECG- and ABP-derived reference measurements was assessed using correlation, concordance, and error metrics. Pearson correlation coefficients (R) were computed to evaluate the strength of the linear relationship between modalities. Lin’s concordance correlation coefficient (CCC) was computed to more directly quantify agreement, as it jointly accounts for correlation and deviation from the identity line. Bland–Altman analysis was performed to assess systematic bias and the limits of agreement (LoA) between the paired measurements. Root mean squared error (RMSE) was computed to characterize overall error magnitude. Linear mixed-effects (LME) modeling was used to account for the differing numbers of data collections per rodent and limited sample size for HRV. The mixed-effects model assessed the relationship between DHI-derived and reference-derived metrics while accounting for repeated observations within subjects through subject-specific random intercepts. The fixed-effect slope of this model represents the proportional relationship between modalities after accounting for subject-level variability, with slopes near unity indicating better agreement. Summary agreement metrics were pooled across data collections to characterize overall DHI system performance for monitoring cardiovascular timing.

## 3. Results

Data were collected from six rats across the cardiovascular monitoring experiments. The adenosine rhythm experiment was performed in one rat with DHI recordings obtained over both exposed vasculature and through intact skin. HRV analysis included eight data collections from five rats, yielding 14 non-overlapping 60-s windows, and PWV analysis in five rats included five data collections and 2756 cardiac cycles.

### 3.1 Heart rate and rhythm characterization

To characterize the ability of our system to record tissue deformation associated with clinically relevant measures of cardiac function, we simultaneously collected DHI and ECG measurements during 60-second trials which included a brief baseline period with normal sinus rhythm followed by rapid intravenous bolus adenosine resulting in atrioventricular (AV) block. These collections were performed both over the exposed femoral artery and through intact skin. The resulting signals are shown in Figure 5.

**Figure 5.**
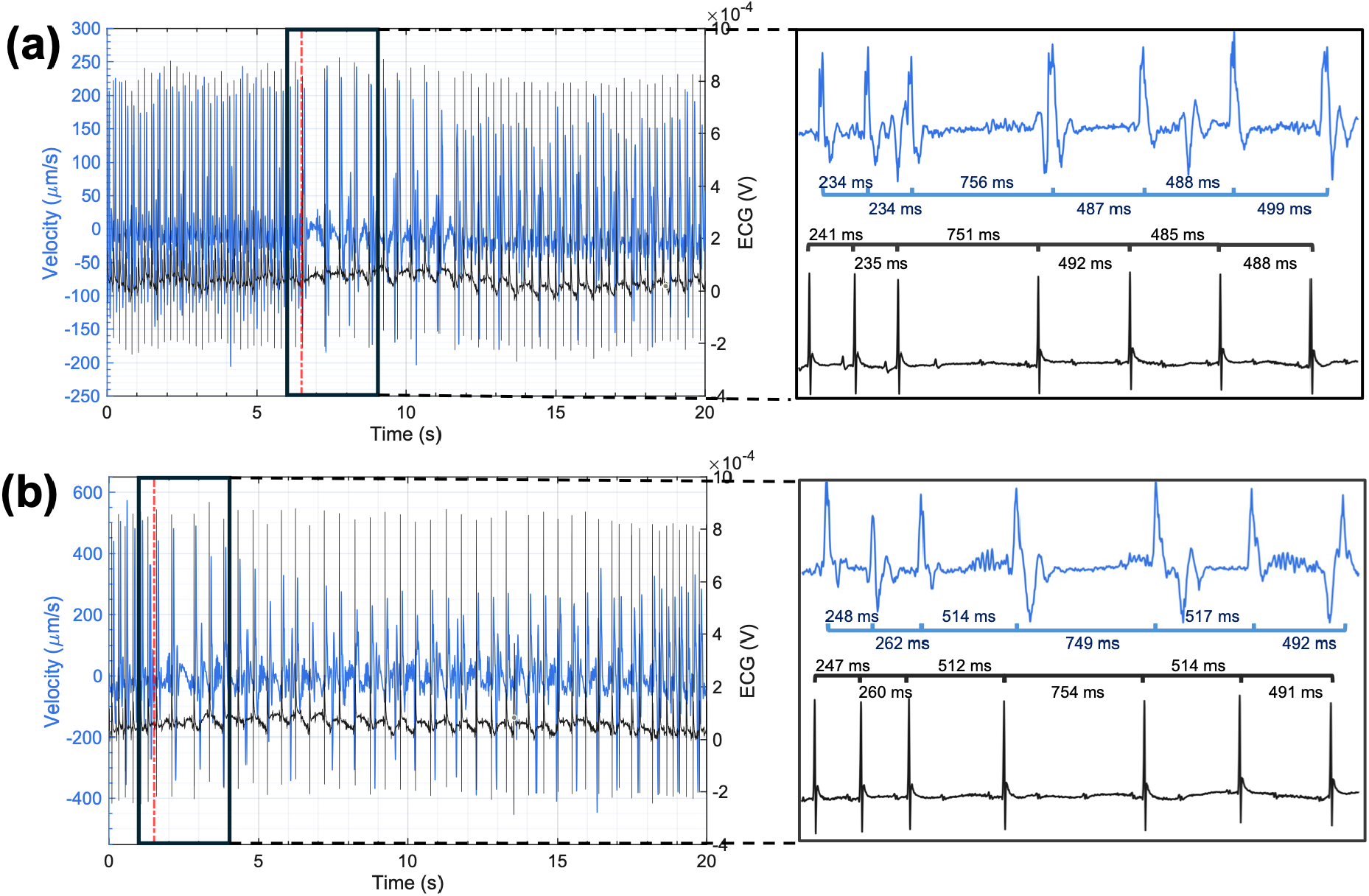
Beat-to-beat cardiac rhythm tracking in tissue velocity and ECG. (a) Synchronously recorded tissue velocity (blue) and ECG (gray) waveforms in the exposed femoral artery during a baseline period followed by administration of intravenous adenosine as indicated by the vertical dashed red line. (b) Synchronously recorded tissue velocity (blue) and ECG (black) waveforms of the femoral artery taken through skin during a baseline period followed by administration of intravenous adenosine indicated by the vertical dashed red line. The insets in (a) and (b) showcase interbeat intervals as tracked by manually identified velocity peaks and ECG peaks in a 3 second interval around adenosine administration.

As demonstrated in the ECG waveform, adenosine altered both cardiac conduction and rate. This was reflected in the slowing of the heart rate shown in Figure 5A and 5B insets. Following administration of adenosine, progression to complete AV heart block occurs and is notable for the absence of the QRS complex. Adenosine is rapidly metabolized with resolution of heart block followed by a shortening of the P-R interval back to baseline and resolution of the bradyarrhythmia to normal sinus rhythm. These cardiac effects can also be seen in the simultaneously recorded tissue velocity waveform from both exposed vasculature and through skin. The time interval between PV features in successive beats closely tracks the ECG RR interval and follows a similar elongation in response to adenosine, followed by a return to baseline level after the adenosine effects washed out.

### 3.2 HRV extraction and characterization

HRV metrics were derived across all 60 s non-overlapping windows from 8 data collections across 5 rodents, resulting in 14 windows (i.e., trials) for analysis. Per window metrics are summarized in Table 2. These results are further visualized in Figure 6.

**Table 2.** DHI- and ECG-derived heart rate and HRV metrics for all trials.

| Trial | Rodent | Mean HR (bpm) |  | SDRR (ms) |  | RMSSD (ms) |  |
| --- | --- | --- | --- | --- | --- | --- | --- |
|  |  | *DHI* | *ECG* | *DHI* | *ECG* | *DHI* | *ECG* |
| 1 | A | 389.0 | 389.0 | 1.6 | 2.0 | 2.0 | 2.6 |
| 2 | B | 308.5 | 308.5 | 4.8 | 3.2 | 7.2 | 4.4 |
| 3 | B | 308.8 | 308.8 | 4.9 | 3.3 | 7.1 | 4.2 |
| 4 | C | 353.9 | 354.2 | 2.1 | 1.9 | 3.2 | 2.2 |
| 5 | C | 349.6 | 349.8 | 2.3 | 1.8 | 3.4 | 1.9 |
| 6 | C | 343.6 | 344.3 | 2.2 | 1.7 | 3.3 | 2.0 |
| 7 | C | 340.6 | 341.2 | 2.4 | 1.6 | 3.6 | 1.9 |
| 8 | D | 334.4 | 334.7 | 4.2 | 3.2 | 6.2 | 4.1 |
| 9 | D | 334.0 | 334.0 | 4.1 | 2.7 | 7.4 | 4.9 |
| 10 | E | 223.0 | 223.0 | 2.5 | 2.1 | 3.6 | 2.4 |
| 11 | E | 224.9 | 224.9 | 2.5 | 2.0 | 3.3 | 2.0 |
| 12 | E | 239.9 | 240.0 | 1.7 | 1.4 | 2.7 | 1.8 |
| 13 | E | 245.6 | 245.7 | 3.5 | 3.3 | 5.7 | 5.1 |
| 14 | E | 244.5 | 244.7 | 3.9 | 3.4 | 5.8 | 5.1 |

**Figure 6.**
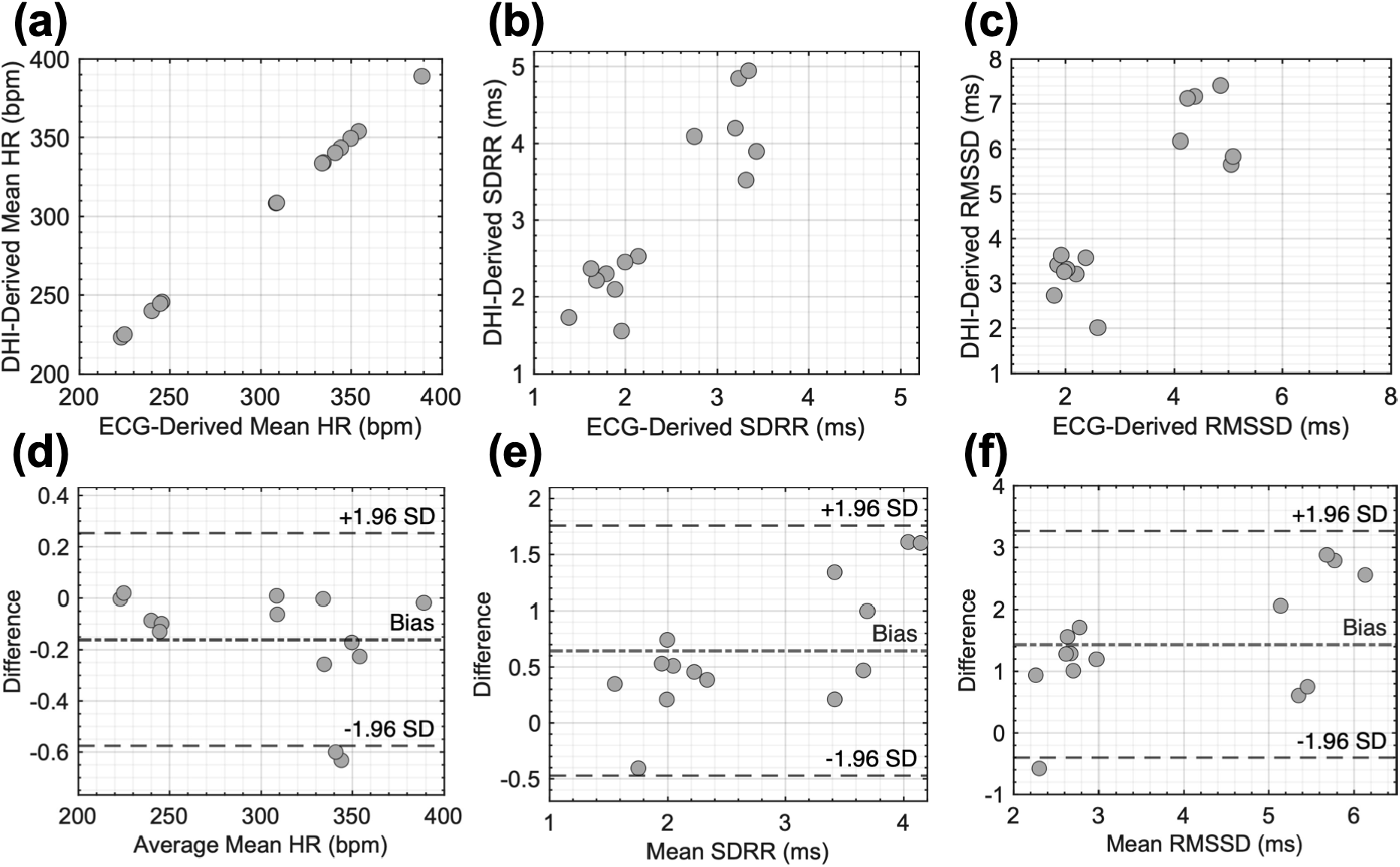
Agreement between ECG-derived and DHI-derived cardiovascular metrics. (A–C) Scatter plots comparing the two modalities for mean heart rate, SDRR, and RMSSD, respectively. (D–F) Corresponding Bland–Altman plots showing the mean bias (central line) and 95% limits of agreement (surrounding dashed lines).

Agreement analysis results between DHI and ECG for these metrics are summarized in Table 3.

**Table 3.**
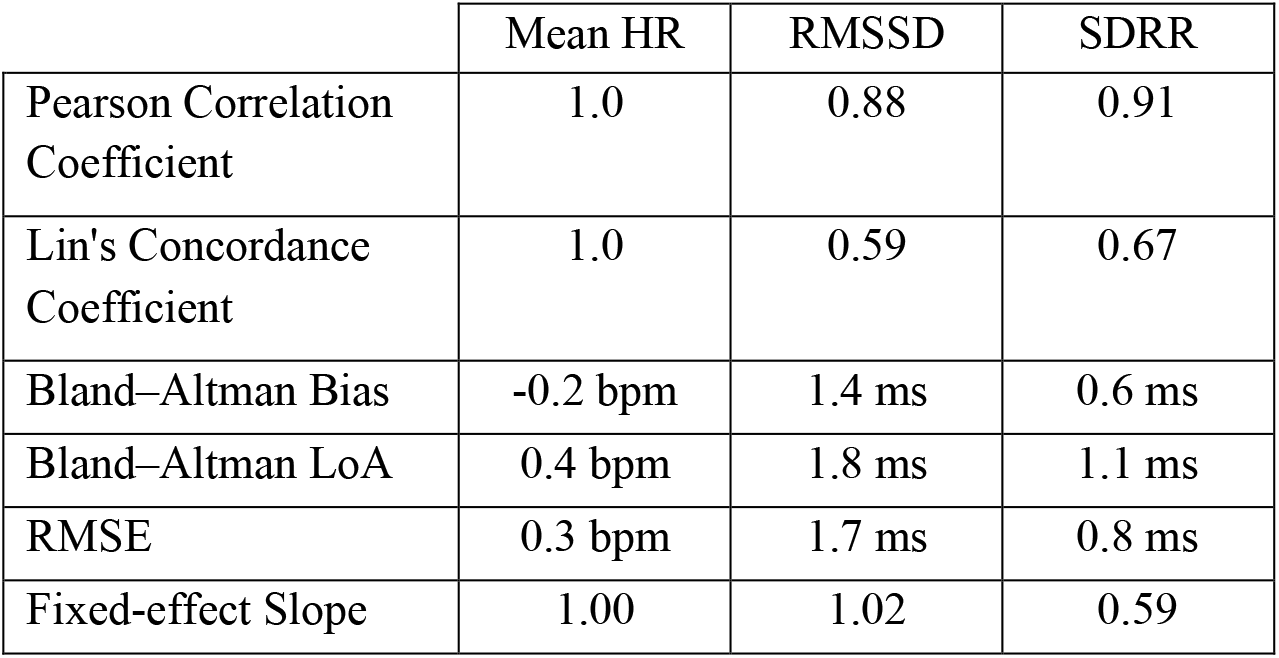
Agreement between DHI- and ECG-derived heart rate and HRV metrics. Values are reported across all paired observations.

The proposed modality demonstrates excellent agreement for mean heart rate estimation and good agreement for HRV metrics in this experimental model. Pearson correlations indicate strong tracking of ECG-derived RMSSD and SDRR values, while CCC values suggest moderate concordance. Bland–Altman analysis indicates a tendency for the proposed modality to overestimate HRV metrics relative to ECG, with moderate limits of agreement observed for RMSSD and SDRR. Mixed-effects modeling further demonstrates significant relationships (α < 0.05) between the modalities after accounting for repeated observations within subjects, supporting the feasibility of the proposed system for proof-of-concept cardiovascular monitoring.

### 3.3 PAT and PWV extraction and characterization

PAT metrics were derived across all cardiac cycles from 5 data collections across 5 rodents. These were then leveraged to derive PWV using the respective sensor’s measured anatomical distance, resulting in 2756 data points for analysis. Per collection PWV results are summarized in Table 4.

**Table 4.** DHI- and ABP-derived average pulse arrival time and pulse wave velocity for all trials.

| Trial | Rodent | d0 + d2 (cm) | d0 + d1 (cm) | # Cycles | Mean PAT (ms) |  | Mean PWV (m/s) |  | PWV RMSE (m/s) |
| --- | --- | --- | --- | --- | --- | --- | --- | --- | --- |
|  |  |  |  |  | DHI | ABP | DHI | ABP |  |
| 1 | A | 11.5 | 10.6 | 384 | 48.9 | 43.6 | 2.35 | 2.43 | 0.08 |
| 2 | B | 11.8 | 11.1 | 613 | 54.3 | 51.4 | 2.18 | 2.16 | 0.07 |
| 3 | F | 7.7 | 6.8 | 494 | 57.0 | 51.5 | 1.35 | 1.32 | 0.04 |
| 4 | D | 10.9 | 10.2 | 789 | 50.9 | 45.8 | 2.14 | 2.23 | 0.09 |
| 5 | E | 13.0 | 11.7 | 476 | 56.7 | 51.8 | 2.30 | 2.26 | 0.06 |

PWV metrics across all collections were pooled for agreement analysis. These results are visualized in Figure 7 and characterized in Table 5.

**Table 5.** Agreement between DHI- and ABP-derived pulse wave velocity. Values are reported across all paired observations.

|  |  |
| --- | --- |
| Pearson Correlation Coefficient | 0.98 |
| Lin's Concordance Coefficient | 0.98 |
| Bland–Altman Bias | -0.02 m/s |
| Bland–Altman LoA | 0.14 m/s |
| RMSE (m/s) | 0.07 m/s |

**Figure 7.**
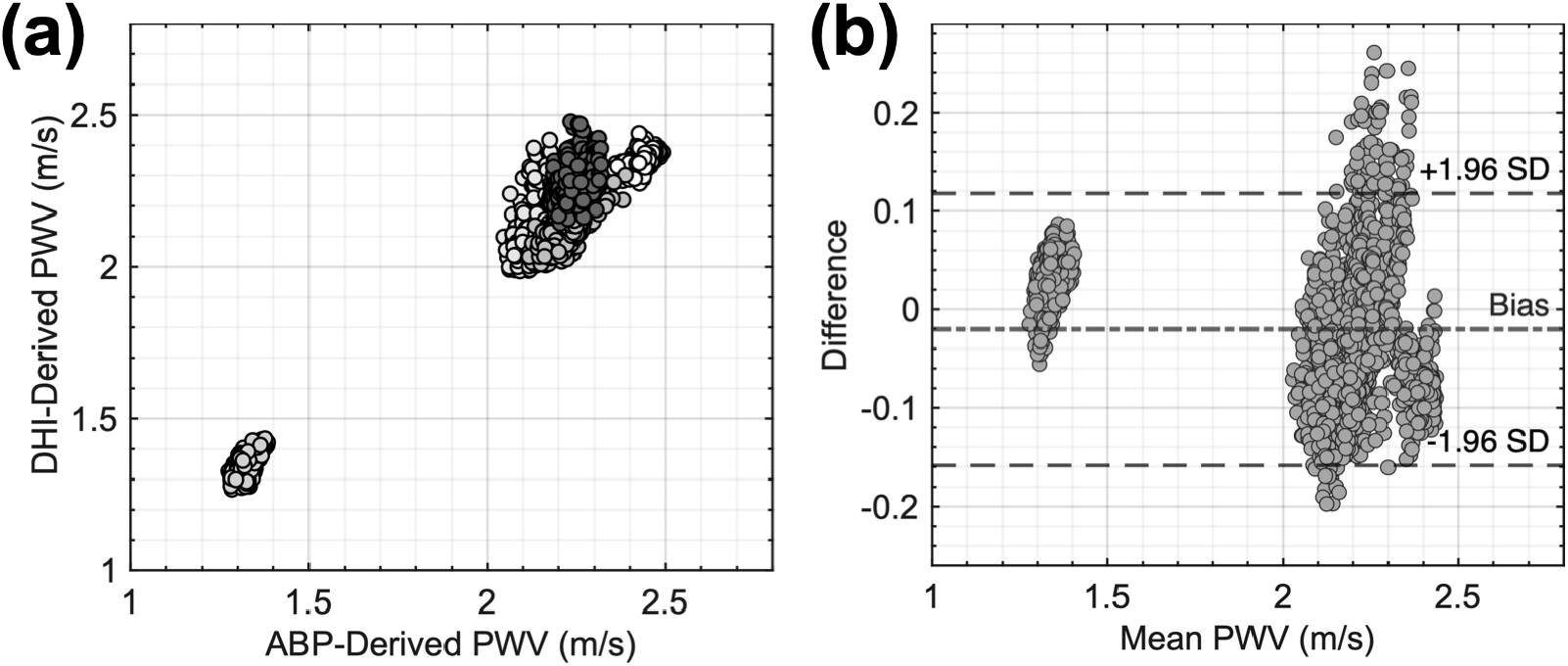
Agreement between ABP-derived and DHI-derived pulse wave velocity. (a) Scatter plot comparing pulse wave velocity derived by pulse arrival recorded by the arterial line (ABP) and by the DHI system. Markers are shade-coded by rodent. (b) Corresponding Bland–Altman plot showing the mean bias (central line) and 95% limits of agreement (surrounding dashed lines).

Agreement analysis results between DHI and the arterial line for PWV are summarized in Table *5*.

The proposed modality demonstrates strong agreement with the reference measurement, with Pearson and Lin’s concordance correlation coefficients of 0.98. Bland–Altman analysis reveals minimal systematic bias (-0.02) and narrow limits of agreement (±0.14), while a low RMSE (0.07) indicates small absolute measurement error. These results suggest that the DHI system produces PWV measurements that closely match the reference method across the evaluated range.

## 4. Discussion

Full-field interferometric measurements of tissue deformation *in vivo* utilizing a DHI system were used to characterize sensitivity to cardiovascular timing, to include heart rate and rhythm as well as measures of pulse propagation time and speed. The tissue velocities described here were predominantly driven by cardiogenic motion as well as respiration, and the waveforms are similar in nature to those from other modalities that record motion such as BCG and SCG [11].

Our system demonstrates high accuracy in tracking physiologically-relevant cardiac timing in these controlled experiments. The sensitivity of DHI extends beyond characterizing heart rate and shows potential to track arrhythmias and beat-to-beat conduction changes. The ECG changes following intravenous adenosine demonstrated the expected progressive AV block (prolongation of the P-R interval) rapidly leading to complete AV block (only P-waves), which was followed by a gradual return to normal sinus rhythm. Velocity waveforms temporally associated with atrial and ventricular contraction demonstrated similar rhythm changes as reflected by the ECG signal on a beat-to-beat basis. Moreover, these changes were able to be tracked in optical measurements taken through skin, indicating the potential for DHI to track cardiovascular information non-invasively. While the data presented are limited, we believe our approach has the potential to non-invasively monitor cardiac arrhythmias, including conduction delays where knowledge of arrhythmias beyond simply heart rate is critical. This condition is particularly relevant in high-risk populations, such as patients following myocardial infarction, in whom high-grade AV block is associated with increased risk of heart failure events, ventricular tachyarrhythmias, and mortality [32,33].

Beyond measuring heart rate and rhythm, our system can also reliably track variability in rhythm via HRV and pulse propagation via PAT and PAT-derived PWV. The ability to record state-dependent indices such as HRV and PAT-derived PWV with a non-contact approach offers exciting potential for a wide range of clinical environments. Characterizing the autonomic nervous system via HRV is increasingly recognized as an important measure for understanding injury severity and prognosis across many clinical domains, including cardiac [34], trauma [35,36], septic [37] and brain-injured patients [38]. In parallel with tracking heart rate and rhythm, as well as indices such as HRV, PWV provides a complementary hemodynamic marker reflecting arterial stiffness and vascular tone, which are strongly influenced by autonomic regulation and cardiovascular status [39,40]. Elevated PWV has been associated with increased cardiovascular risk [41], impaired vascular compliance [39], and hypertension [42,43]. Simultaneous continuous recording of these indices via DHI could support improved clinical decision making.

Tracking tissue velocity with a full-field interferometric imaging system allows for spatial aggregation across many pixels and flexible signal formation, providing advantages over single-point measurements such as LDV. Spatial aggregation across multiple spatial channels provides signal diversity across partially independent speckle realizations, reducing speckle-induced noise and signal dropout relative to a single-point measurement [44]. In addition, imaging enables spatially informed processing, such that signals can be formed over anatomically relevant ROIs while excluding noisy areas without repositioning the sensor. This full-field access also increases the physiological information that could be extracted from a single recording, as distinct dynamics from different tissue types and structures may be contained within the field of view. This spatial information could be leveraged to probe regional tissue function. Moreover, the interferometric basis of DHI produces calibrated nanometer-scale displacements at high temporal bandwidths, which may provide meaningful improvements over intensity-based imaging systems and ultimately enable sensitivity to faster mechanical events. There are many possible avenues of future investigation to better understand the information within tissue motion at high spatiotemporal resolutions.

There were several experimental and processing decisions that may have impacted the results. In particular, selection of the frame rate impacts spatial resolution, as higher frame rates necessitate smaller FPAs in the camera used in this study. There may be a tradeoff in signal quality between temporal bandwidth and the number of pixels available for spatial aggregation, which was not explored in this study. As has been demonstrated from other non-contact approaches for vital sign monitoring, tuning camera parameters is critical to measurement of various physiologic waveforms [45]. Another consideration is that data was collected over the femoral artery, which may not be the optimal anatomical location for tracking heart rate variability. As has been shown by other modalities, sensor location impacts the morphology and timing of tissue motion recorded [46,47]. A limitation of this study is that most of the data analyzed were recorded from exposed vasculature, and further validation is needed to verify that these results are maintained when imaging through intact skin to enable non-invasive monitoring. Moreover, the data presented in this study were collected over relatively short time scales, and verifying stability over longer collection durations will likely be important. Lastly, and perhaps most importantly, it remains to be determined whether this approach will generalize to humans. We anticipate that our system sensitivity should be adequate for measuring cardiogenic motion in humans; however, uncertainty remains regarding noise and artifacts, and these must be understood and accounted for before assessing performance in a clinical environment.

Overall, this work should be interpreted as a feasibility study demonstrating that DHI can recover multiple cardiovascular timing metrics from a single non-contact measurement in controlled rodent preparation. The results of our study represent an initial step in demonstrating the viability of full-field interferometry in non-contact recording of cardiac physiology, and future work is needed to validate performance through intact skin, over longer durations, and in the presence of realistic motion and noise in human subjects.

## 5. Conclusion

Recording tissue deformation at a high spatiotemporal resolution with a non-contact DHI system, we demonstrate the ability to continuously track normal and abnormal heart rate and rhythm as well as prolonged state-dependent changes via heart rate variability and pulse wave velocity. Non-contact full-field interferometric imaging could therefore enable real-time monitoring of clinically relevant physiologic changes and is a promising tool to ultimately aid clinicians in the treatment of patients.

## Funding

This work was supported by Johns Hopkins University Discovery Award funding and by Johns Hopkins Applied Physics Laboratory internal research and development funding.

## Acknowledgments

ChatGPT-5 was used in editing for language clarity and grammar.

## Disclosures

ATL, NES, CLR, EB, and DWB are named inventors on U.S. Patent Nos. 11,733,650 and 11,625,003, which are related to the technology described in this work. The remaining authors declare no potential conflicts of interest with respect to the research, authorship, or publication of this article.

## Code, Data, and Materials availability

Data in support of the findings of this paper are available upon reasonable request.

